# The Distribution of Enset Pests and Pathogens and a Genomic Survey of Enset Xanthomonas Wilt

**DOI:** 10.1101/2020.06.18.144261

**Authors:** Zerihun Yemataw, James S. Borrell, Manosh Kumar Biswas, Oliver White, Wendawek Mengesha, Sadik Muzemil, Jaypal N. Darbar, Ian Ondo, Pat J.S. Heslop Harrison, Guy Blomme, Paul Wilkin

**Affiliations:** Southern Agricultural Research Institute, Hawassa, Southern Nations Nationalities and Peoples Regional State, Ethiopia; Department of Natural Capital and Plant Health, Royal Botanic Gardens, Kew, Richmond, Surrey, TW9 3AE, UK; Department of Genetics and Genome Biology, University of Leicester, LR1 7RH, UK; Department of Biology, Hawassa University, Hawassa, Ethiopia; School of Life Sciences, University of Warwick, Coventry CV4 7AL, UK; Bioversity International, Addis Ababa office, c/o ILRI, P.O. Box 5689, Addis Ababa, Ethiopia

**Keywords:** Bacterial wilt, Enset Xanthomonas Wilt, food security, plant health, root mealybug, Xanthomonas Wilt of enset

## Abstract

Mapping the distribution of crop pests and pathogens is essential to safeguard food security and sustainable livelihoods. However, these data are unavailable for many neglected and underutilised crops, particularly in developing countries. In Ethiopia, the world’s largest historic recipient of food aid, the indigenous banana relative enset (*Ensete ventricosum*) is threatened by multiple pests and pathogens whilst providing the staple starch source for 20 million people. Foremost among these is *Xanthomonas* Wilt of enset (EXW), caused by *Xanthomonas vasicola* pv. *musacearum* (*Xvm*), a globally important disease of bananas (*Musa* sp.) that likely originated in enset. Here we collate 1069 farm surveys to map the distribution and relative prevalence of enset pests and pathogens across the entire enset growing region. We find that EXW is the most frequently encountered pathogen, and that farmers consistently ranked EXW as the most significant constraint on enset agriculture. Our surveys also showed that corm rot, and the pests root mealybug, mole rat and porcupine are all virtually ubiquitous. Finally, we apply genotyping-by-sequencing to the detection of *Xvm* and demonstrate that it is present even in asymptomatic domesticated and wild enset samples, suggesting that management of plants displaying symptoms alone may not be sufficient to reduce disease transmission. Holistic understanding of pests and pathogen distributions in enset may have significant benefits for both food security in Ethiopia, and preventing proliferation in related crops such as banana across central and east Africa.

## Introduction

The increasing transmission of plant pests and pathogens has significant consequences for the distribution, quality and yield of crops (Bebber et al. 2014; Savary et al. 2019). Rural subsistence farmers appear particularly susceptible to these impacts, where emergence or outbreaks of pests and pathogens exacerbates existing food insecurity (Bruce 2010; Vurro et al. 2010) or hinders agricultural resilience (Heeb et al. 2019). Whilst global surveillance systems exist for pest and pathogens of major crops (Forum et al. 2019), basic distribution, prevalence and incidence data is missing for many neglected and underutilized plants which are likely to become increasingly important in future diversified food systems (Borrell et al. 2019).

This paucity of monitoring data is a major challenge in Ethiopia where enset (*Ensete ventricosum* (Welw.) Cheesman), an indigenous banana relative, provides food security for 20 million people, but is threatened by multiple poorly documented pests and pathogens (Jones 2000, 2018; Blomme *et al*. 2017; Borrell *et al*. 2019). Enset cultivation is largely restricted to south and southwest Ethiopia (Figure 1A) where it is grown principally as a subsistence crop and for regional markets, and often comprises a significant proportion of total farm area (Borrell et al. 2020; Sahle et al. 2018). Enset is a monocarpic perennial that can grow for up to a decade before reaching maturity and is readily vegetatively propagated. Farmers maintain a cycle of plantings and transplantings of various ages that can be harvested at any time prior to flowering and senescence. Following harvest, the pseudostem and corm are pulped and fermented to provide a storable starch source (Tamrat et al. 2020). This flexible system enables farmers to buffer seasonal food deficits, earning enset the moniker ‘the tree against hunger’ (Brandt et al. 1997).

**Figure 1.**
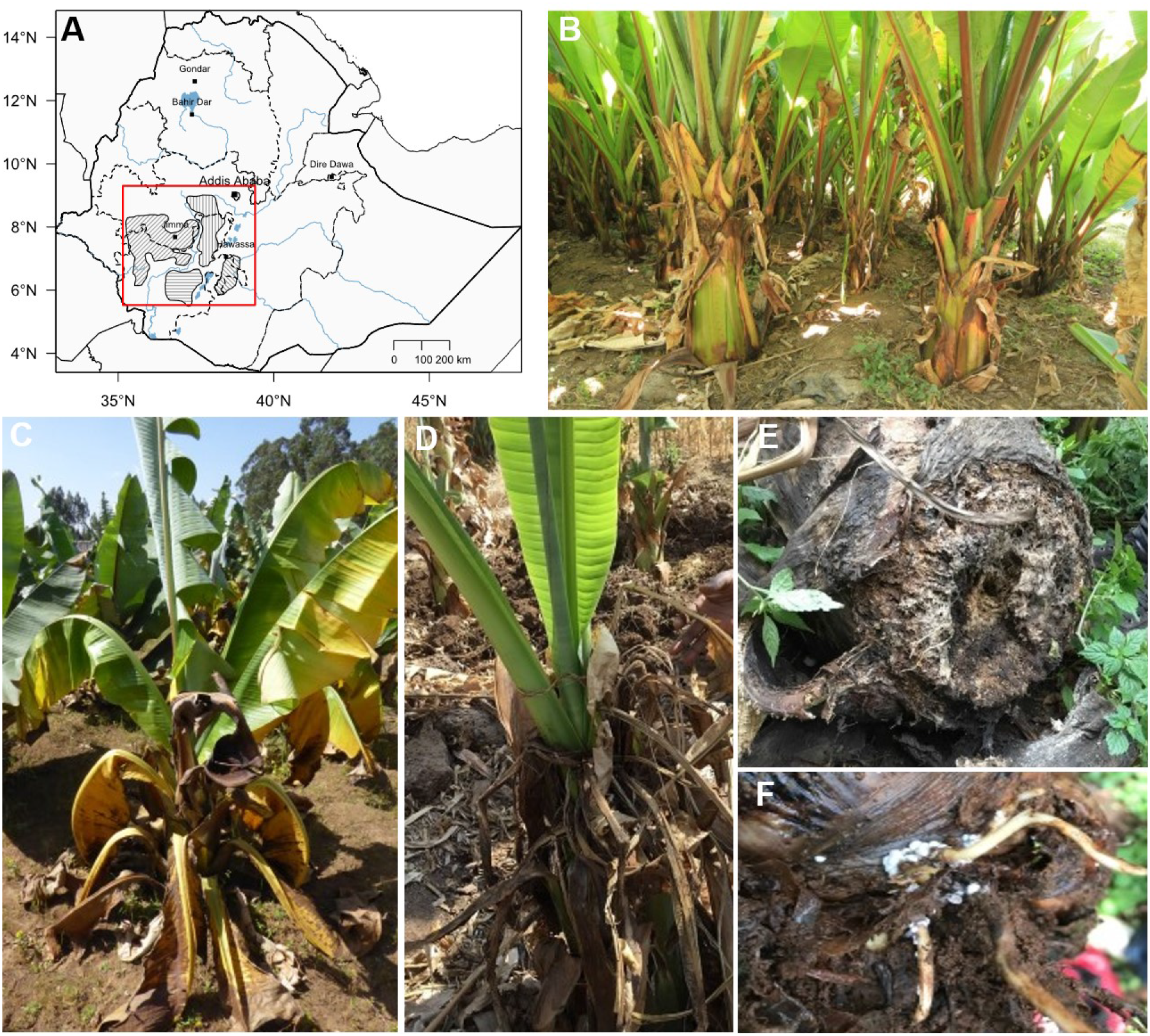
Study area and major enset pests and pathogens in Ethiopia. A) Map of Ethiopia, with shaded polygons denoting main regions of enset agriculture and red boundary indicating the extent of our study area. B) Typical enset plot. C) Enset with symptoms of Enset Xanthomonas Wilt (EXW). D) Enset of landrace ‘Badadet’ apparently recovering from severe EXW. E) Enset with evidence of corm rot. F) Root mealybugs on enset corm and roots.

Enset production is affected by multiple pests and pathogens of varying severity (Table 1, Figure 1). Important biotic constraints include Enset *Xanthomonas* Wilt (EXW, bacterial rots (*Erwinia* sp.) and root mealybugs (Blomme et al. 2017; Bogale et al. 2004; Addis, Azerefegne, Blomme, et al. 2008; Tewodros and Tesfaye 2014; Shank and Ertiro 1996). EXW is caused by *Xanthomonas vasicola* pv. *musacearum (Xvm*) (formerly *X. campestris* pv. *musacearum*). For clarity, in this manuscript we follow Studholme et al., 2019 and refer to the causal organism as *X. vasicola* pv. *musacearum*, except when discussing NCBI the reference genome which is still accessioned as *X. campestris* pv. *musacearum*. Mammal pests that directly damage the plants include porcupine, mole rats, wild pigs and monkeys (Bobosha 2003), and these are also suspected vectors of disease transmission, especially EXW, hence they are included in this study (Hunduma et al. 2015; Pers. Obs. J.S. Borrell).

**Table 1.**
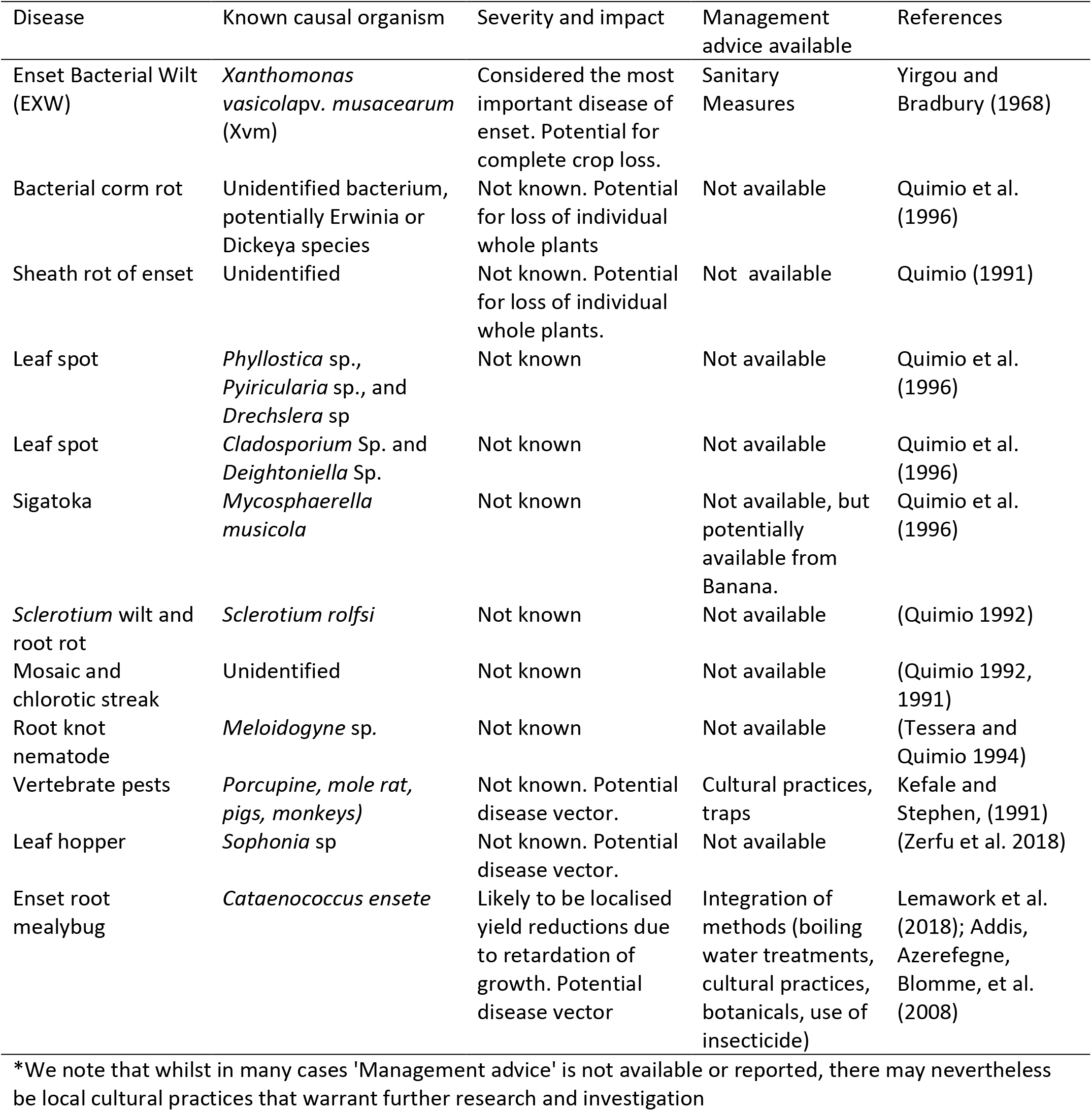
Reported pests and pathogens of domesticated enset (*Ensete ventricosum*) in Ethiopia.

Among these pests and pathogens, EXW is frequently cited as the most significant concern for farmers, generating a large number of studies that seek to identify tolerant or resistant enset landraces (Hunduma et al. 2015; G Welde-Michael et al. 2008; Yemataw et al. 2016; Muzemil et al. 2020; Haile et al. 2020) *Xvm* infects the vascular system of enset, damaging the harvestable tissue, causing permanent wilting and eventually death (Yemataw et al. 2017). It is known to be transmitted by contaminated tools and infected planting material, and potentially biotic vectors, such as wild and domesticated animals that browse part of the corm or pseudostem (Yemataw et al. 2017; Addis et al. 2010). In a previous study across 320 farms in eight districts, 40% of respondents reported EXW in their field (McKnight-CCRP 2013), though this varied by region from 3.3% (Kembata Tembaro) to 95.7% (Gedeo), and the authors suggest that the true infection rate (farm prevalence) could be as high as 80%. Some authors claim that EXW has forced farmers to abandon enset production (Spring 1996; Tadesse et al. 2003). EXW is also speculated to be a possible driver for a reported major historic decline in enset agriculture in the North of Ethiopia (∼200 years b.p.), though there is a lack of evidence to support or refute this (Brandt et al. 1997).

The causative agent of EXW was first described by Yirgou and Bradbury (1968) in Ethiopia. However symptoms consistent with EXW were reported as early as the 1930s (Castellani 1939; Studholme et al. 2019; Blomme et al. 2017), though it is not clear whether this represents emergence of the disease, or simply the first scientific documentation. During the 1960-80s the pathogen spread rapidly in enset and banana (*Musa* sp.) in Ethiopia (Yemataw et al. 2017) and is now a threat to smallholder banana cultivation throughout central and eastern Africa (Carter et al. 2010), impacting food security and rural livelihoods (Blomme et al. 2013, 2017). As a result, improved understanding of *Xvm’s* spatial distribution, intensity and impact on farmers is key to continued food security, as well as supporting translational research in enset and bananas (Merga et al. 2019). Two previous studies surveyed banana *Xanthomonas* wilt (BXW) in the East African highlands (not including Ethiopia) (Bouwmeester et al. 2016) and the risk of BXW more widely across Africa (Ocimati et al. 2019), but not at a resolution that is informative for disease mapping or management in Ethiopia, the putative origin of the disease.

Compared with *Xvm*/EXW, the distribution, prevalence and impact of other enset pests and pathogens has received much less attention. Enset corm rot is thought to be caused by *Erwinia* (or *Dickeya*) species (Blomme et al. 2017), but is poorly characterised. A survey by Yirgu (2016) in the Gamo Highlands found that a quarter of respondents considered corm rot to be the most severe disease of enset. Enset root mealybug (*Cataenococcus ensete* Williams and Matile-Ferrero) is known to be a locally important pest, with evidence that infestation retards growth, reduces pseudostem circumference and associated yields (Addis *et al*. 2008; Azerefegne *et al*. 2009). Addis (2005) reported that 30% of sampled farms were infected. Limited surveys of nematodes and weevils were undertaken by Bogale *et al*. (2004), which found relatively low nematode densities and did not find weevils. The banana weevil *Cosmopolites sordidus* does not thrive well above 1,600 m asl (Lescot 1988), and most enset cultivation zones are located at higher altitudes. Leaf hopper was found to be widespread in Yem special district, and associated with EXW prevalence (Zerfu et al. 2018). There remains the possibility of additional undescribed pathogens, in both wild and domesticated populations.

Here, we apply spatial and molecular methods to undertake the most extensive survey to date of the pests and pathogens affecting enset agriculture in Ethiopia, with a particular focus on detecting EXW. To achieve this, we first use region-wide farmer interviews and farm surveys to evaluate the relative abundance of pests and pathogens on enset farms, and farmer perceptions of the major constraints on enset agriculture. Second, we collate a suite of high-resolution environmental, topographic and socioeconomic variables for the study area and apply these to characterise the spatial distribution and prevalence of major enset pests and pathogens across the enset growing region. Finally, we apply a genotyping-by-sequence approach to survey the leaf-associated microbiota of EXW-symptomatic and non-symptomatic enset samples, to assess detection efficacy of diseased versus incubating or asymptomatic *Xvm* and improve our understanding of EXW transmission. We discuss these data in the context of ongoing monitoring of pests and pathogens in a neglected food security crop, to support diagnosis, monitoring and management.

## Materials and Methods

### Enset pests and pathogen surveys

This study comprises observations from two region-wide surveys, conducted independently by i) the Southern Agricultural Research Institute between 2014-17 (n=585), hereafter SARI and ii) a team from Royal Botanic Gardens Kew, Wolkite and Hawassa Universities 2017-20 (n=484), hereafter KWH. Both surveys were independently conceived, designed and carried out over a broadly similar geographic area (Figure 2A), using a similar methodology and as a result we are confident that comparing and combining these data make our conclusions more robust.

**Figure 2.**
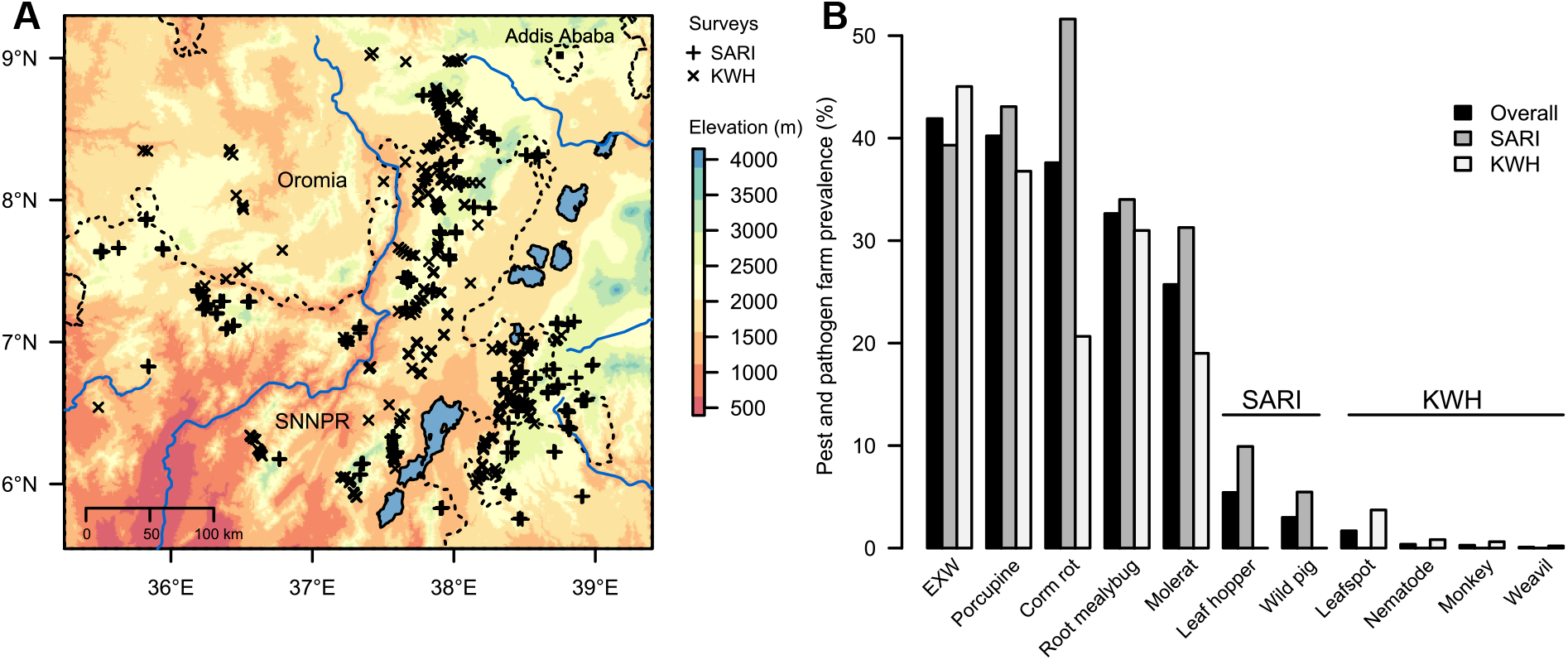
Summary of enset pest and pathogen surveys. A) Spatial distribution of the two independent enset pest and pathogen surveys analysed in this study. B) Percentage of farm surveys that recorded each of 11 enset pests and pathogens.

Both surveys were conducted by experienced teams together with local agricultural extension agents, and randomly selected individual farms over stratified sampling regions. Data was collected via individual interviews and direct on-farm participatory observations, and all diseases scored as presence or absence. Both surveys recorded the presence of five major pests and pathogens: EXW, root mealybug, corm rot, mole rat and porcupine, as well as estimating the number of enset plants with symptoms of EXW. The SARI survey additionally recorded leaf hoppers and wild pig damage, and asked farmers to identify the most important constraint on enset production. The KWH survey additionally recorded nematodes, leafspot, monkey damage and weevils. Survey methodologies were broadly consistent; applying semi-structured interview, disease identification sheets and visual inspection of infected or damaged plants. The order in which diseases were presented, and the diagnostic photographs shown were varied to mitigate reporting bias. We also sought to document any symptoms of similar magnitude that were not attributed to known pests and diseases.

We calculated the relative prevalence of pests and pathogens to compare consistency across surveys. Where a pest or pathogen was only recorded in one survey, we record prevalence relative to the number of farms in that survey. Due to road accessibility and logistics, surveys were conducted unevenly through the year. Therefore, these data are not sufficiently robust for assessing seasonal trends, though we provide a summary of seasonality in Supplementary Materials (Figure S1). Finally, we grouped farms based on the top five farmer perceived constraints on enset agriculture and compared these with pests and pathogens prevalence.

### Spatial modelling of pest and pathogen prevalence

We collated 41 high-resolution environmental, topographic and socioeconomic variables. Environmental variables were sourced from WorldClim (Fick and Hijmans 2017), ENVIREM (Title and Bemmels 2018) and CliMond (Kriticos et al. 2012), together with a 90m SRTM DEM sourced from Jarvis et al. (2008). Slope, aspect, topographic position index and terrain roughness were calculated from elevation using the ‘terrain’ function in the R package Raster (Hijmans 2017). Socioeconomic variables were derived from OpenStreetMap (OpenStreetMap contributors 2015) and Gridded Population of the World v4 (Center for International Earth Science Information Network 2017). All variables were resampled to 250m resolution, for consistency with the high-resolution topographic variables. A full list of variables is provided in Supplementary Materials, Table S1. Despite the resilience of our chosen analysis approach to highly correlated variables, to aid subsequent interpretation we removed 26 variables with very high collinearity using the function vifcor in ‘usdm’ (th=0.8) (Naimi 2017; Lever et al. 2017). All analyses were conducted in R software (R Core team 2019).

To build robust models, we tested a range of cluster aggregation values to group farms by distance and sample size (see validation section), selecting a maximum aggregation distance of 4000m and ≥ 3 surveyed farms (Supplementary Materials, Figure S2). We chose a relatively fine scale aggregation due to the high environmental heterogeneity of southwest Ethiopia. Prevalence was calculated as the proportion of farms affected within a cluster. Observation clusters with <3 surveyed farms were excluded from model building. Environmental variables were then extracted for each surveyed farm, and averaged by cluster.

To characterise the climatic niche of each enset pest and pathogen we used an approach similar to that of Pironon *et al*. (2019). First, principal component analysis (PCA) was performed on 100,000 systematically sampled points representing background climatic space of the study area. In these analyses, the first two principal components summarise the variation of the 15 retained variables. Second, we computed quantiles of pest and pathogen prevalence from our survey data corresponding to the 10^th^, 50^th^ and 90^th^ percentile. To characterise the niche occupied by a given pest or pathogen we plotted an alpha hull for each degree of pest and pathogen severity using the package ‘alphahull’ with an alpha value of 1.05 (Pateiro-lopez 2019). This approach visualises the most severely affected farm clusters nested within the broader climatic space of surveyed farms. Climatic space polygons were then replotted in geographical space using the R packages ‘raster’ and ‘rgeos’ (Hijmans 2017; Bivand et al. 2018).

To provide an indication of the strength of association between environmental variables and our predicted pest and pathogen niches, we randomly sampled each variable across our four overlapping prevalence polygons (0, 0.1, 0.5 and 0.9) and estimated the Kendall rank correlation coefficient. We note that we did not estimate the significance of each association, as this would be strongly influenced by the number of random samples. Finally, we used the nicheOverlap function in the R package Dismo (Hijmans et al. 2017) to estimate niche overlap between modelled pest and pathogen distributions for 0.1 and 0.9 prevalence quantiles.

### Model validation

We used three approaches to validate our spatial analysis. First, we performed a sensitivity analysis by varying the aggregation distance and cluster threshold size of farm surveys, then assessing the change in predicted area as a response. Second, we modelled data from each survey separately and evaluated performance by comparing predicted area. Finally, for EXW we use a generalized linear model to test the hypothesis that aggregated survey points with greater disease prevalence also display greater disease severity in the number of infected plants.

### Diseased tissue sample collection and genotyping

We collected leaf tissue samples from 10 enset individuals (multiple landraces), displaying EXW symptoms. This was complemented by 233 domesticated and 14 wild enset samples that did not display visible symptoms and were otherwise considered healthy. Samples were widely distributed across the study area, with a maximum of three from a single farm. Samples were principally collected for diversity analysis meaning they encompass a broad range of putatively genetically distinct landraces. Leaf tissue was silica dried, extracted using a standard CTAB protocol (Doyle & Doyle 1987), normalised and submitted to Data2Bio (IA, USA) for library preparation and tunable genotyping-by-sequencing (tGBS) following the protocol of Ott *et al*. (2017). DNA samples were digested with the restriction enzymes NspI and BfcCI/Sau3AI before being sequenced using an Ion Proton platform.

### Identification of candidate bacterial sequences

We screened all samples for putative bacterial sequences by implementing a local blast search (Camacho et al. 2009) against a custom database of bacterial genome sequences created using NCBI Reference Sequences (RefSeq; O’Leary et al. 2016). Specifically, we downloaded all complete bacterial genomes classified as “reference” or “representative” resulting in a dataset comprised of 3,000 assemblies (date accessed 12^th^ June 2020; Supplementary Table 2). In addition, we included genome sequences for *Xanthomonas campestris* pv. *musacearum* (GenBank accession: GCA_000277875.1) and *Xanthomonas vasicola* pv. *vasculorum* (GCA_003015715.1). *Xanthomonas campestris* pv. *musacearum* was used as it is the causal agent of bacterial wilt in enset and banana which has recently been reclassified as *X. vasicola* pv. *musacearum* (Aritua et al. 2008). *Xanthomonas vasicola* pv. *vasculorum* was included as it is a close relative of *Xvm* yet is non-pathogenic in banana (Wasukira et al. 2012).

Prior to the blast search, raw tGBS sequencing reads were quality filtered using Trimmomatic (Bolger et al. 2014). Duplicate sequences were filtered from our samples using CD-HIT (Fu et al. 2012). A blastn search of tGBS reads was performed against the custom bacterial genome refseq dataset with 10 maximum target sequences, one maximum high-scoring segment pair (HSP) and an expectation value (E) of 1 ×10^−25^. The taxonomy of a query sequence was defined using a “best sum bitscore” approach, where the bitscores for each subject taxonomy identified are summed and the taxonomy with the greatest score is selected. Where more than one taxonomy has the greatest sum bitscore no taxonomy is defined. This avoids ambiguous assignment with multiple closely related taxa in the blast database. Our approach was adapted from the methodology of blobtools2 (Challis et al. 2020) which was not appropriate for our analyses as it does not distinguish subspecific taxonomic ranks (i.e. pathovars).

For an overview of the bacteria present in and/or on leaf tissue, we first counted sequences assigned to each genus or species in our blast dataset. Taxa were scored as present in an individual if we identified >5 matching reads for that sample. To provide a sequence-depth independent estimate, we then calculated the base pair coverage of the *X. vasicola* pv. *musacearum* genome that was identified in our blast search. To do this, overlapping blast hits for *X. vasicola* pv. *musacearum* in each sample were merged using bedr in R (R Core team 2019) and total base pair coverage was calculated. We plotted these data in putative groupings comprising diseased, non-diseased and wild samples, and applied Analysis of Variance (ANOVA) and Tukey HSD post-hoc tests to assess differences between groups. Custom scripts used for the blast search, taxonomic identification and coverage estimation are available form https://github.com/o-william-white/Enset_tGBS.

## Results

### Farm and farmer surveys

A total of 1069 farms were assessed across two survey campaigns (Figure 2A). Overall, EXW was the most frequently recorded pest or pathogen, occurring in 41.2% of farms with porcupine (40.2%) and corm rot (37.6%) also similarly abundant (Figure 2B). In a comparison between surveys, corm rot, followed by porcupine and EXW was most frequently encountered by SARI, whereas EXW, followed by porcupine and root mealybug was most frequently encountered by HWK. Whilst the study area was largely consistent, the distribution of survey effort across months differed between surveys, with the majority of SARI survey effort in November-December and HWK in October and January-April (Supplementary Materials, Figure S1). Of 577 farmer responses, 507 (88%) reported pests and pathogens as the predominant constraint on enset agriculture (Table 2). Of the remainder, 34 respondents reported no major constraint and others cited eight additional abiotic constraints at low frequency, including drought, land shortage, frost and labour shortage. Farmer perception of the predominant constraint on enset agriculture was highly consistent with the frequency at which pest and pathogens were recorded on farms (Table 2).

**Table 2.** Comparison of farmer perceived constraints and associated proportion of farms on which each pest or pathogen was recorded. Columns one and two denote the reported constraint; subsequent columns show the proportion of those farms in which each pest or pathogen was recorded.

| Farmer reported constraint | Number of reports* | Proportion of farms in which species was recorded |  |  |  |  |
| --- | --- | --- | --- | --- | --- | --- |
|  |  | EXW | Root mealybug | Corm rot | Molerat | Porcupine |
| EXW | 265 (45.9%) | 0.71 | 0.29 | 0.48 | 0.31 | 0.42 |
| Root mealybug | 48 (8.3%) | 0.15 | 0.98 | 0.31 | 0.13 | 0.38 |
| Corm rot | 87 (15.1%) | 0.07 | 0.21 | 1.00 | 0.13 | 0.28 |
| Molerat | 49 (8.5%) | 0.20 | 0.45 | 0.39 | 0.98 | 0.47 |
| Porcupine | 53 (9.2%) | 0.17 | 0.23 | 0.47 | 0.26 | 0.98 |

\* Three additional respondents reported Wild pigs, one leaf blight and one leaf hopper.

### Spatial modelling of pests and pathogens

We computed the niche space for the five major pest and pathogen species with sufficient data and projected these into geographical space (Figure 3). Estimation of Kendall rank correlation coefficients between environmental variables and modelled pest and pathogen prevalence quantiles identified different suites of variables for each species (Table 3). EXW was positively associated continentality and negatively associated with the maximum temperature of the coldest month and potential evapotranspiration (PET) of the coldest quarter. Corm rot was negatively associated with multiple PET variables and root mealybug was negatively associated with isothermality and cold quarter precipitation. Pairwise niche overlap at the 0.9 prevalence quantile was highest for EXW – porcupine and EXW – corm rot respectively (Table 4).

**Figure 3.**
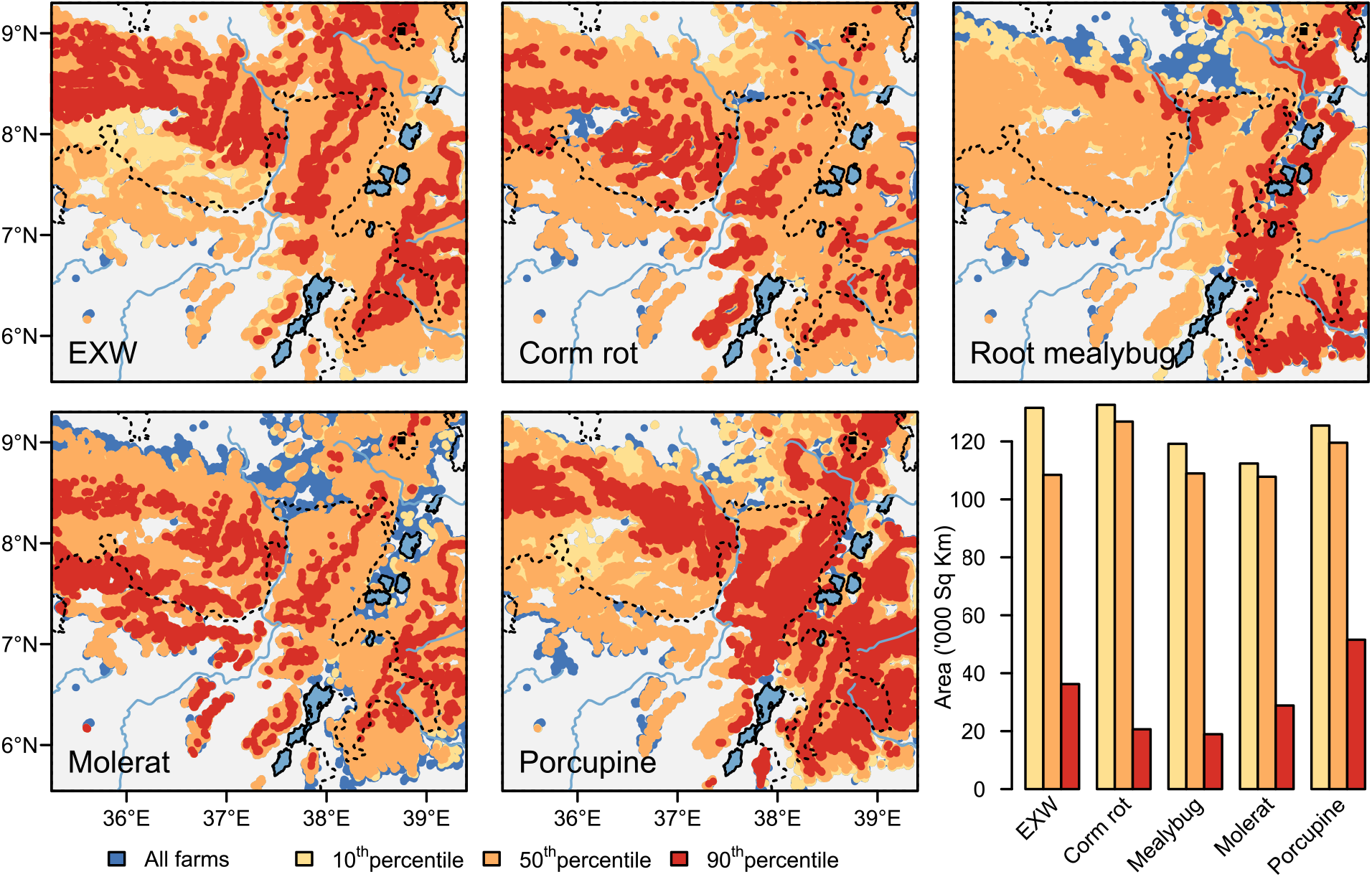
Distribution maps of five major enset pests and pathogens. Colour scales depict quantiles of pest and pathogen prevalence in farm survey clusters. Barchart depicts the area (‘000 km^2^) of pest and pathogen occurrence at each prevalence quantile.

**Table 3.** Kendall rank correlation coefficient for each spatial variable and the PCA-derived distribution of each pest and pathogen.

| Spatial variables | EXW | Corm<br>rot | Root<br>mealybug | Molerat | Porcupine |
| --- | --- | --- | --- | --- | --- |
| Thornthwaite aridity index | 0.038 | -0.039 | -0.010 | -0.090 | 0.007 |
| Precipitation of wettest week | 0.014 | 0.020 | -0.055 | 0.022 | -0.091 |
| Precipitation of warmest quarter | -0.008 | 0.027 | -0.088 | 0.097 | -0.026 |
| Precipitation of coldest quarter | 0.016 | 0.030 | -0.129 | 0.035 | -0.078 |
| Mean diurnal temperature range | 0.030 | -0.022 | 0.008 | -0.091 | 0.050 |
| Isothermality | -0.037 | 0.042 | -0.122 | 0.039 | -0.036 |
| Climatic Moisture Index | 0.001 | 0.044 | -0.076 | 0.090 | -0.102 |
| continentality | 0.050 | -0.055 | 0.060 | -0.104 | 0.017 |
| Max Temp. Coldest | -0.047 | -0.053 | 0.048 | -0.054 | 0.044 |
| PET Coldest Quarter | -0.050 | -0.046 | 0.118 | -0.008 | 0.060 |
| PET Driest Quarter | -0.010 | -0.044 | 0.124 | -0.075 | 0.054 |
| PET Wettest Quarter | 0.030 | -0.080 | 0.068 | -0.105 | 0.021 |
| Topographic position Index | 0.004 | -0.009 | -0.009 | -0.014 | 0.012 |
| Elevation | 0.032 | 0.039 | -0.030 | 0.048 | -0.029 |
| Distance to a major road | 0.001 | 0.045 | 0.008 | 0.038 | -0.047 |

**Table 4.** Pairwise niche overlap across the five major enset pests and pathogens. Lower triangle indicates overlap at 10% prevalence quantile. Upper triangle indicates overlap at 90% prevalence quantile.

| Pests and pathogens | EXW | Corm rot | Root mealybug | Molerat | Porcupine |
| --- | --- | --- | --- | --- | --- |
| EXW | - | 0.32 | 0.01 | 0.05 | 0.33 |
| Corm rot | 0.96 | - | 0.02 | 0.04 | 0.24 |
| Root mealybug | 0.91 | 0.91 | - | 0.00 | 0.20 |
| Molerat | 0.84 | 0.85 | 0.91 | - | 0.03 |
| Porcupine | 0.96 | 0.96 | 0.94 | 0.87 | - |

### Validation

We plotted the predicted area for each of the five species (at 0.5 and 0.9 disease prevalence quantiles) across a range of cluster aggregation distances and cluster minimum sizes (Supplementary Materials, Figure S3). These show that predicted areas for most species stabilize at 4000m and clusters of 3 or more farms. At very high cluster values (>7), the predicted area declined, which we attribute to declining sample sizes reducing our ability to predict across climatic space. Comparison of models derived from each survey individually showed a significant correlation in predicted area (F_1,13_ = 115, R^2^ = 0.89, p = <0.001) (Supplementary Materials, Figure S4). Finally, we report a highly significant relationship between the proportion of farm clusters reporting EXW and the mean count of EXW infected enset (F_118_ = 1.78, p = 0.008) (Figure S5).

### Genetic survey of EXW

In a large sample of visibly non-diseased domesticated plants, bacteria of the genera *Acinetobacter, Cyanobacterium*, and *Pseudomonas* were most abundant, whilst *Nostoc* and *Oscillatoria* were less abundant but recorded in a high proportion of individuals (Figure 4A). A similar, but more diverse and abundant assemblage was recorded in visibly non-diseased wild plants, including the genera *Methylobacterium* and *Sphingomonas* (Figure 4C). We note that whilst *Methylobacterium* has been reported as a frequent laboratory contaminant (Salter et al. 2014), here it is largely localised to wild samples extracted using multiple kits and sequenced on different plates, suggesting this is a valid finding. By contrast, the diseased domesticated sample group was characterised by a very high mean number of reads of *Xanthomonas*, present in all samples (Figure 4B). It is noteworthy, however, that reads corresponding to *Xanthomonas* were also identified in >57% of non-diseased domesticated samples and >86% of wild samples. Of the 19 *Xanthomonas* reference genomes in our blast database, *X. campestris* pv. *musacearum* (i.e. *Xvm*) was the most frequently identified species (Figure 4D). Finally, retaining only reads aligning to *Xvm*, we calculated the coverage of the blast hits against the genome for each sample and plotted these by group (Figure 4E). The recovered sequence length significantly differed across groups (F_(2,254)_=46.2, P=<0.001), despite no significant difference in raw read counts between the three groups (F_(2,254)_ = 1.08, p = 0.34). A Tukey post-hoc test indicated significant *Xvm* genome coverage differences between the domesticated and diseased groups (p < 0.001) and wild and diseased groups (p < 0.001). However, there was no significant difference between wild and domesticated (p = 0.99). Of 233 asymptomatic domesticated plant samples, 36 (15%) reported a count of *Xvm* aligning reads equal to or higher than an EXW symptomatic sample. A further 150 (64%) plants reported a non-zero number of *Xvm* aligning reads. We plot the distribution of our diseased reference plants and the 36 asymptomatic plants with an equal, or higher number of *Xvm* reads in Figure 5.

**Figure 4.**
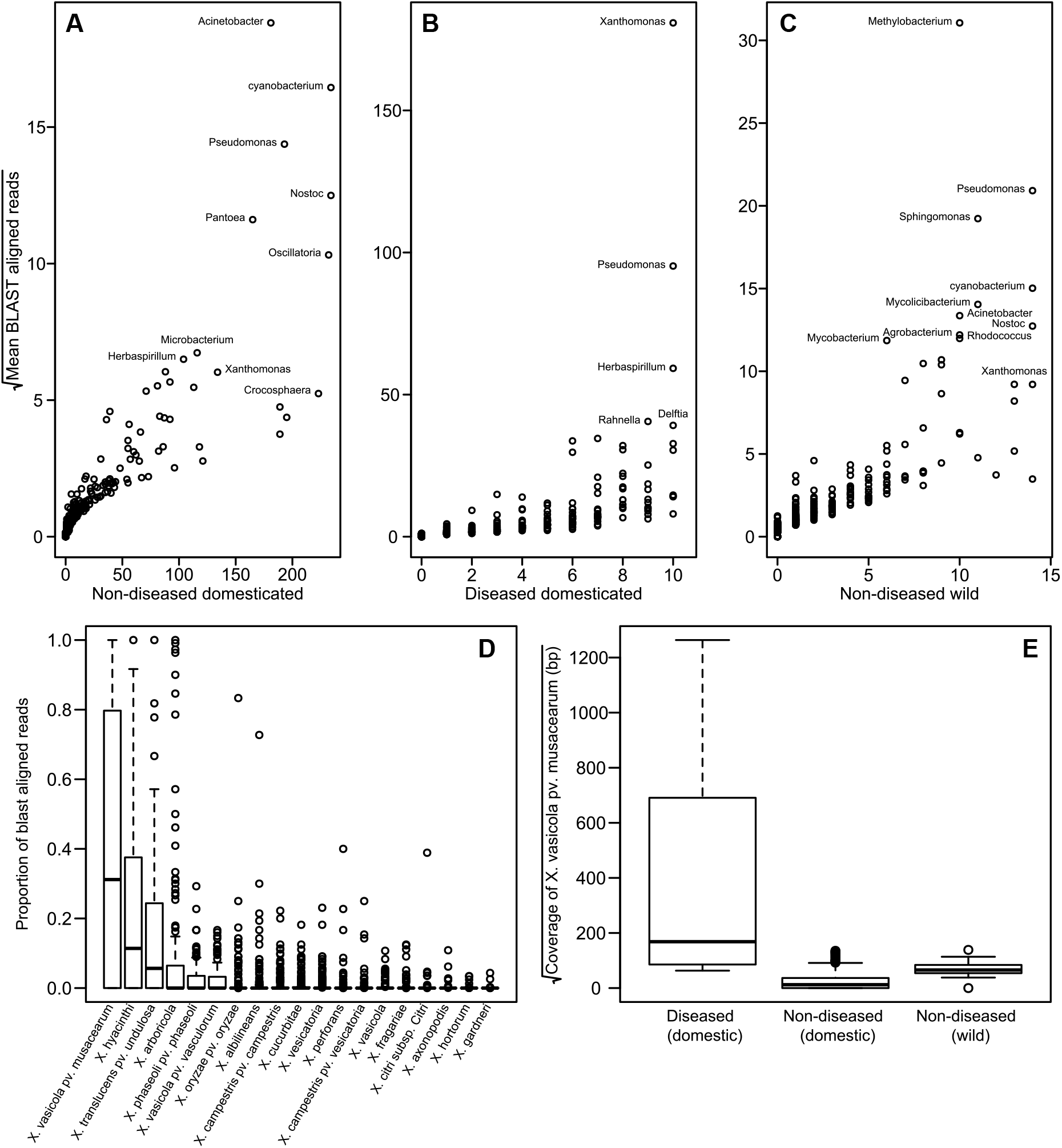
Enset leaf microbiome characterisation based on blast aligned raw genotyping-by-sequencing reads. A-C) Microbial genera identified in diseased and non-diseased enset samples. D) Number of reads aligning to each of 19 *Xanthomonas* reference sequences. E) Total coverage of *Xanthomonas vasicola* pv. *musacearum* blast aligned reads for each enset sample, grouped by disease status and domestication.

**Figure 5.**
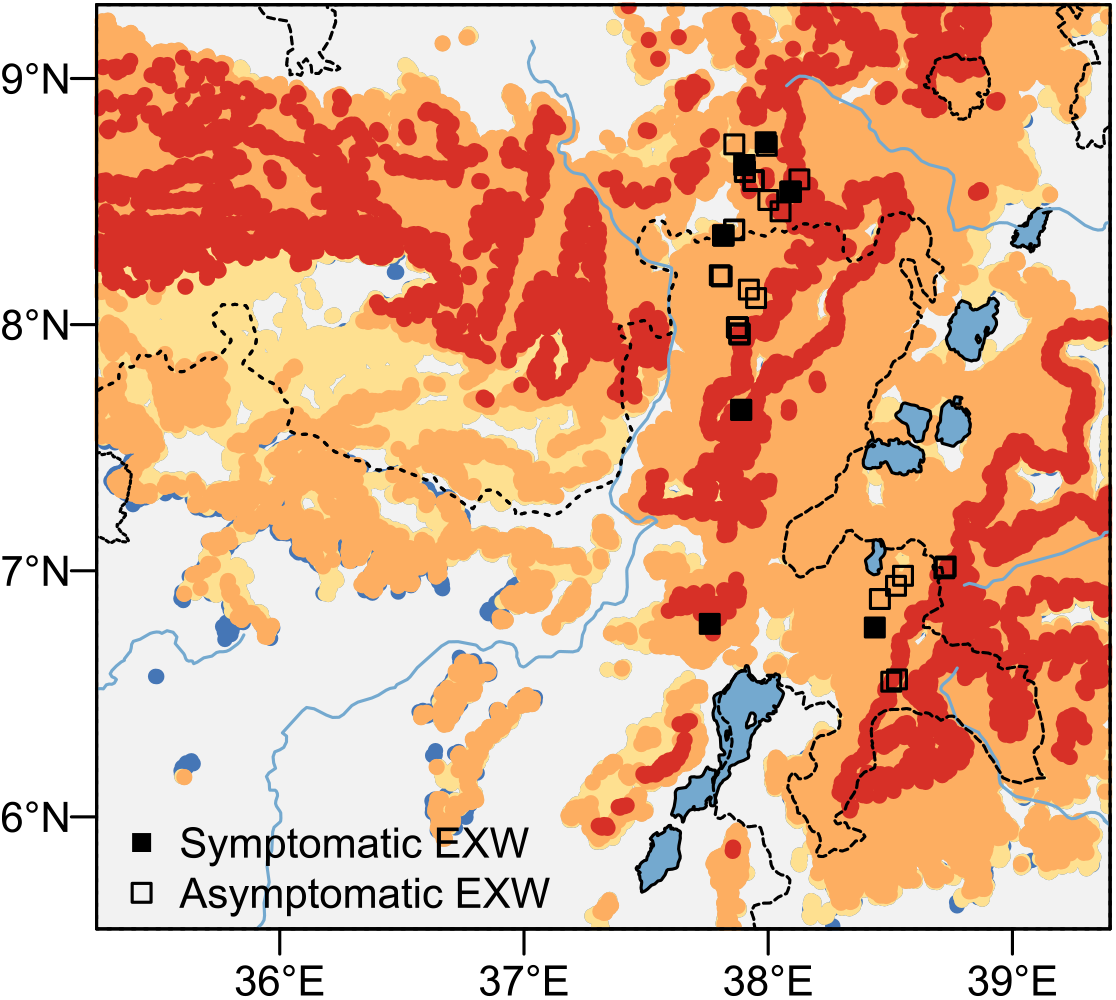
Distribution of EXW symptomatic and asymptomatic enset samples in south west Ethiopia. Background map is modelled enset distribution (Figure 3). The minimum *Xvm* read count in symptomatic samples was 4000. Here, we plot the 37 asymptomatic individuals with an equal or greater number of *Xvm* reads.

## Discussion

The distribution and intensity of pests and pathogens in neglected and underutilised crop species is often poorly known, limiting the effectiveness of mitigation strategies (Bebber et al. 2019). In this study we collate 1069 farm surveys to provide the most detailed analysis to date of the distribution and prevalence of pest and pathogens on enset farms in Ethiopia, together with the perceptions of enset farmers, to develop a baseline from which to assess future trends.

### Farmer surveys

The most frequently recorded pest or pathogen was EXW, occurring on 42% of farms (Figure 2). This is consistent with multiple previous reports emphasising the importance of EXW as a constraint to enset agriculture (Shimelash et al. 2008; Merga et al. 2019; Wolde et al. 2016). However, we found that other much less studied pests and pathogens are also virtually ubiquitous across the enset growing region. Corm rot and root mealybug were reported on 37.6% and 32.6 % of farms respectively. Whilst we know that both EXW and corm rot can result in loss of whole plants, the associated reduction in yield due to mealybug infestation is not known.

Despite large sample sizes, we observed differences in disease prevalence between the two surveys. These may be attributed to a number of causes, particularly survey timing and variation of disease prevalence across overlapping survey areas. Though our sampling strategy was not designed to ascertain seasonal trends (and so must be used with caution for this purpose), we note that SARI performed a large number of surveys in November and December, a period shortly after the long rainy season in which we see an increase in the number of corm rot observations (Supplementary materials, Figure S1). This may partly account for the disparity in corm rot observations. Observations for EXW, root mealybug and porcupine were largely consistent between surveys. These comparisons emphasise the value of multiple independent surveys, particularly where pathogens are poorly known and may be cryptic.

Our findings are consistent with farmer perceptions, with over 40% reporting EXW to be the primary constraint on enset agriculture (Table 2). However, only 71% of farmers that report EXW as the primary constraint were found to have EXW on their farms. This is in contrast with farmers who reported other pests and pathogens as their primary constraint, where our surveys found the reported pest or pathogen present on the farm >97% of the time. Therefore, EXW is considered a greater constraint by farmers than would be assumed from its frequency alone. This could be due to the potentially devastating impact of EXW, and the risk of greater livelihood and food security consequences than from other pests and pathogens (Azerefegne *et al*. 2009; Savary *et al*. 2012; Borrell *et al*. 2019). For banana farmers in East and Central Africa, Banana Xanthomonas Wilt (BXW) is also ranked above other pests and pathogens (Tushemereirwe et al. 2006; Blomme et al. 2017).

The prevalence of EXW on Ethiopian enset farms is similar or slightly less severe than published reports of BXW elsewhere in east Africa. In a large survey in Uganda, Nakato *et al*. (2016) recorded BXW in 69-75% of farms, and in Rwanda, Uwamahoro et al. (2019) found prevalence varied from 26-82% of farms. If EXW is indeed less prevalent, it is possible that a) enset is less susceptible to *Xvm*, b) its presence for nearly a century has helped farmers select for a larger proportion of tolerant landraces, c) the lack of insect-vectored transmission with enset reduces observed prevalence and especially incidence and/or d) another factor such as environment or cultural practice reduces disease prevalence. Relatively few surveys in Ethiopia have focused on BXW, to provide a comparison, though a study by Shimelash *et al*. (2008) reported that the number of infected plants varied from ∼2-40% across a series of sampling sites stratified by elevation. Major banana growing regions in Ethiopia (e.g. Arba Minch) are largely spatially and altitudinally isolated from the principal area of enset cultivation, which may have served to limit the incidence of *Xvm* in lowland banana production zones.

### Distribution of pest and pathogens

We modelled the distribution of five major pest and pathogens and found all to be virtually ubiquitous across the survey area (Figure 3). This helps explain why previous studies have found it challenging to identify hotspots in pathogens such as EXW (Wolde et al. 2016; Brandt et al. 1997). Our observations were largely consistent across a range of parameters and both independent surveys (Figures S3, S4). Despite the broad distribution of most pests and pathogens, we did observe variation in relative disease prevalence consistent with our limited knowledge on pest and pathogen ecology. For example, the most severely affected regions for Root mealybug appear to be low lying areas along the Great Rift Valley, consistent with reports that mealybugs are most common in moist, humid localities and that they can be dispersed via flooding events (Azerefegne et al. 2009). Similarly, we show that corm rot is negatively correlated with drier localities (higher potential evapotranspiration) which are likely to be less amenable to bacterial multiplication. EXW was more weakly associated with multiple environmental variables, the most important being maximum temperatures and potential evapotranspiration in the coldest quarter. Interestingly, EXW was most strongly positively associated with Continentality (average temperature of the warmest month, minus coldest month). Higher Continentality values are typical of areas where domesticated has expanded beyond the range of wild enset.

Additional unmeasured variables are also likely to be important in refining these models and our understanding of enset disease ecology. For example, the wide diversity of cultural practices may regionally facilitate or hinder control of pests or transmission of pathogens. Whilst we did not find that ‘distance to roads’ was a strong predictor as might have been expected for pathogens such as EXW and root mealybug that can be transmitted through planting materials, other socioeconomic variables such as farm density or the proportion of enset in the local crop mix, may be important. Whilst some data on the prevalence of enset agriculture is available (Borrell et al. 2020), these data are not at a sufficiently high resolution to be analytically tractable. Pests and pathogens may also vary in their ecology and virulence (Goss and Bergelson 2006). For example, researchers screening enset landraces for *Xvm* tolerance have reported varying virulence across *Xvm* isolates (Muzemil et al. 2020; Merga et al. 2019; G Welde-Michael et al. 2008). Finally, while our data captures pest and pathogen farm-level prevalence, it does not quantify disease incidence or severity i.e. the impact on yield or livelihoods. Future surveys focused on quantifying yield reduction would complement this work.

### Evidence of EXW in asymptomatic plants

In this study, we are confident that we have detected *X. vasicola* pv. *musacearum* in ‘diseased’ samples as they display the known disease phenotype and a large number of reads blast align to the *Xvm* genome sequence (Figure 4). Surprisingly, we also detected a significant number of *Xvm* reads in asymptomatic domesticated and wild enset plant samples. There are three possible explanations for this observation. First, a subset of landraces may display some tolerance or resistance meaning that the pathogen can be present without causing symptoms. Second, we may be detecting a non-pathogenic or closely related *Xanthomonas* pathovar in our asymptomatic samples (Alemayehu et al. 2016). This is supported by the fact that variation in the pathogenicity of different strains has been reported previously (G Welde-Michael et al. 2008; Merga et al. 2019; Muzemil et al. 2020; Haile et al. 2020). Finally, it is possible that we are detecting *Xvm* during the incubation period. The incubation period in enset appears to be longer than in *Musa*, though this depends on the infected landrace, entry point of the pathogen, inoculum level and age of the plants (Ocimati et al. 2013; G. Welde-Michael et al. 2008; G Welde-Michael et al. 2008). In this case we would conclude that *Xvm* is present, and may eventually cause symptoms. We note that long term latent infections have been reported in East african bananas (Ocimati et al. 2013). It is also noteworthy that we detect *Xvm* in wild enset, which is consistent with reports by Alemayehu *et al*. (2016) of wild enset susceptibility. However unlike Alemayehu *et al*., throughout our extensive fieldwork we have not observed a wild enset plant displaying EXW symptoms and it is not clear how *Xvm* could cross generations in a wild unmanaged population.

We consider it plausible that all three explanations may be responsible to varying degrees. Therefore, whilst we have likely detected *Xvm* during incubation in some individuals, this probably does not explain detection of *Xvm* in nearly half of asymptomatic domesticated plants and nearly all wild plants. Therefore, tolerance of low levels of *Xvm* and the existence of latent infections, coupled with a possible wider diversity and varying pathogenicity of *Xvm* in enset agricultural systems, suggests that the overall distribution of this pathogen may have been underestimated.

### Current research gaps

Building on the first region wide pest and pathogen distribution maps, we attempt to identify major outstanding research gaps. Firstly, whilst a growing number of studies are surveying putatively EXW tolerant or resistant enset landraces it remains to be understood why EXW appears to predominantly affect 4-5 year old plants (Wolde et al. 2016) (or whether that is simply observation bias as these are likely to be the most common group demographically). Secondly, abiotic stress prior to infection can predispose plants to pathogen susceptibility (Bostock et al. 2014). It is possible that susceptibility to EXW is exacerbated by abiotic stress, such as drought or cold shock, though the underlying processes may be much more complex (Neil et al. 2017). It would be worthwhile to identify a stratified sample of farms for continued repeat EXW surveys to understand seasonality patterns in severity. Similarly, transmission may be higher under certain environmental conditions. Shimwela *et al*. (2016) reported higher BXW incidence during the rainy season, attributed to higher water levels in plant tissue favouring bacteria development. This suggests that transmission can also be higher in wet conditions as inoculum levels may be elevated (Blomme et al. 2017), and tool use increases for management reasons. Finally, we have not addressed potential interactions, for example, whether root mealybug infestation makes enset more susceptible to EXW, or facilitates entry of the pathogen into the roots or corm. However we note the strong niche overlap between porcupine (as a putative vector) and both EXW and corm rot (Table 4).

### Conclusions

In conclusion, farmers clearly consider EXW to be the predominant constraint on enset agriculture. Their concern may be justified based on evidence presented here that *Xvm* is more widespread and prevalent than previously recognised, partly explaining the propensity of EXW to appear unexpectedly. In a regional context, *Xvm* it can be considered one of the most important and widespread disease of *Musa* in East and Central Africa with significant economic and food security impacts. Whilst EXW has proven to be a substantial challenge for effective disease management in small scale farming settings, comparatively less research has been undertaken on corm rot and root mealybug, which our data demonstrates are similarly widespread and prevalent. Whilst they may not have the potential severity of EXW, they may cumulatively have a significant impact on overall yields and food security. Despite the significant challenges that pathogens such as *Xvm* pose, enset agriculture is rich in indigenous knowledge, genetically diverse landraces and a wide range of agronomic practices; significantly more so than in the introduced (in Ethiopia) genetically depauperate and agronomically uniform *Musa* crop, which predominantly focusses on the Cavendish dessert banana types. This suggests that further research in enset may have translational benefits for related species in Ethiopia and beyond.

## Acknowledgements

We thank the Southern Agricultural Research Institute for providing logistical support, and numerous agricultural extension agents for facilitating fieldwork.

## Author Contributions

ZY, JB, WM and SM performed field surveys and collated data. JB designed and performed spatial analysis with contributions from IO. MB and JD sequenced enset tissue samples and OW processed and analysed sequence data. JB wrote the first draft of the manuscript and produced the figures. All authors contributed to and approved the final version of the manuscript.

