## Supplementary Materials for "The Distribution of Enset Pests and Pathogens and a Genomic Survey of Enset Xanthomonas Wilt"

### Supplementary Tables

**Table S1.** Environmental, topographic and socioeconomic variables used in pest and pathogen niche modelling.

| Variable | Description | Units |
|---|---|---|
| Bio01 | Annual mean temperature | °C |
| Bio02 | Mean diurnal temperature range | °C |
| Bio03 | Isothermality (Bio02 ÷ Bio07) | - |
| Bio04 | Temperature seasonality | CV |
| Bio05 | Max temperature of warmest week | °C |
| Bio06 | Min temperature of coldest week | °C |
| Bio07 | Temperature annual range (Bio05-Bio06) | °C |
| Bio08 | Mean temperature of wettest quarter | °C |
| Bio09 | Mean temperature of driest quarter | °C |
| Bio10 | Mean temperature of warmest quarter | °C |
| Bio11 | Mean temperature of coldest quarter | °C |
| Bio12 | Annual precipitation | mm |
| Bio13 | Precipitation of wettest week | mm |
| Bio14 | Precipitation of driest week | mm |
| Bio15 | Precipitation seasonality | CV |
| Bio16 | Precipitation of wettest quarter | mm |
| Bio17 | Precipitation of driest quarter | mm |
| Bio18 | Precipitation of warmest quarter | mm |
| Bio19 | Precipitation of coldest quarter | mm |
| Annual PET | annual potential evapotranspiration | mm/year |
| Thornthwaite aridity index | Index of the degree of water deficit below water need | - |
| Climatic Moisture Index | a metric of relative wetness and aridity | - |
| continentality | average temp. of warmest month - average temp. of coldest month | °C |
| Emberger'S Q | Emberger's pluviothermic quotient: a metric that was designed to differentiate among Mediterranean type climates | - |
| Growing Degree Days (0) | Sum of mean monthly temperature for months with mean temperature greater than 0°C multiplied by number of days | - |

| | | |
|---|---|---|
| Max Temp. Coldest | Maximum temp of the coldest month | ℃ * 10 |
| minTempWarmest | Minimum temp of the warmest month | ℃ * 10 |
| PET Coldest Quarter | Mean monthly Potential Evapotranspiration of the Coldest Quarter | mm / month |
| PET Driest Quarter | Mean monthly Potential Evapotranspiration of the Driest Quarter | mm / month |
| PET Warmest Quarter | Mean monthly Potential Evapotranspiration of the warmest Quarter | mm / month |
| PET Wettest Quarter | Mean monthly Potential Evapotranspiration of the wettest Quarter | mm / month |
| Thermicity Index | Sum of mean annual temp., min. temp. of coldest month, max. temp. of the coldest month, x 10 | °C |
| Slope | Mean elevation difference between focal cell and eight neighbouring cells | m |
| Aspect | The compass direction that a slope faces in degrees from North | ° |
| Topographic position Index | Mean of the absolute difference between the value of a cell and the eight surrounding cells | m |
| Roughness | Largest inter-cell difference between a central cell and the eight neighbouring cells | m |
| Elevation | meters above sea level | m |
| Road distance | Distance from grid cell centroid to nearest road segment. | Km |
| Line Density | Number of road segments crossing each cell | - |
| Travel time | Accessibility map expressed as the travel time in minutes to different-sized cities | min |
| Population density | Population density (by district) | p / km2 |

**Table S2.** Bacterial reference genome dataset used for blast analysis originating from NCBI Reference Sequences and created using RefSeq (O'Leary et al. 2016) (date accessed 12th June 2020).

[see separate file]

### Supplementary Figures

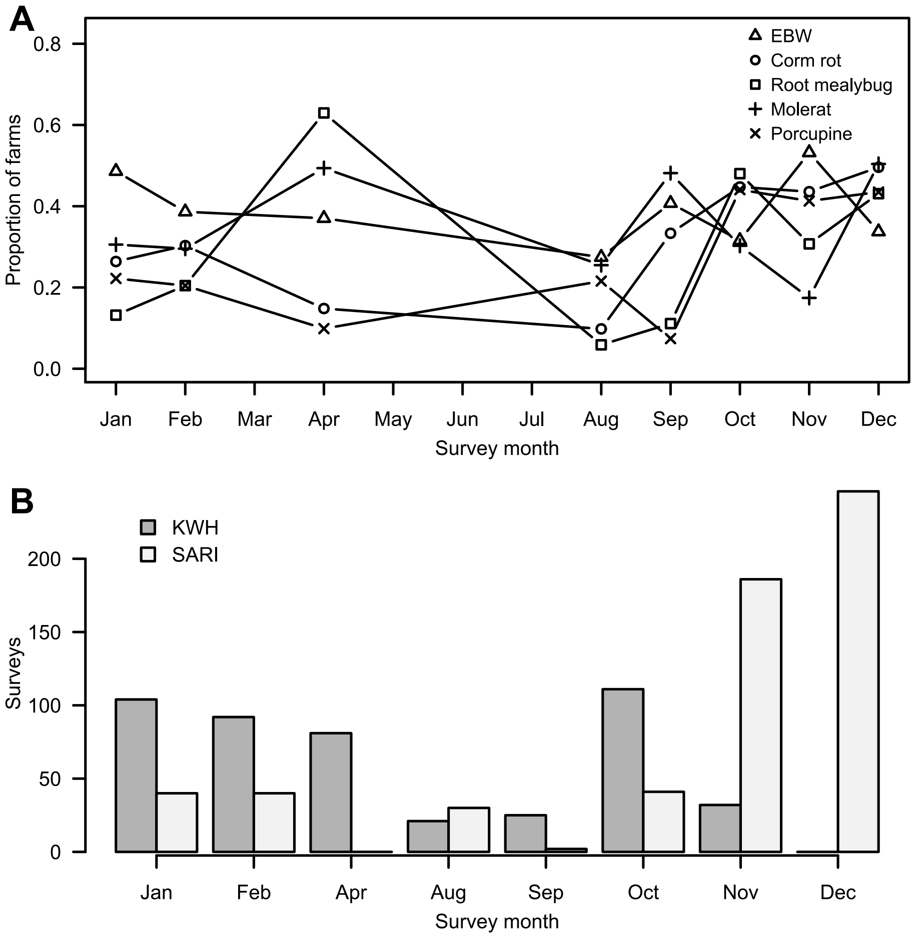


**Figure S1. Distribution of enset farm surveys and pest and pathogen records over time.** A) Seasonal variation in proportion of farms with observed enset pest and pathogens. B) Comparison of survey timing. Note surveys were also conducted across discrete years.

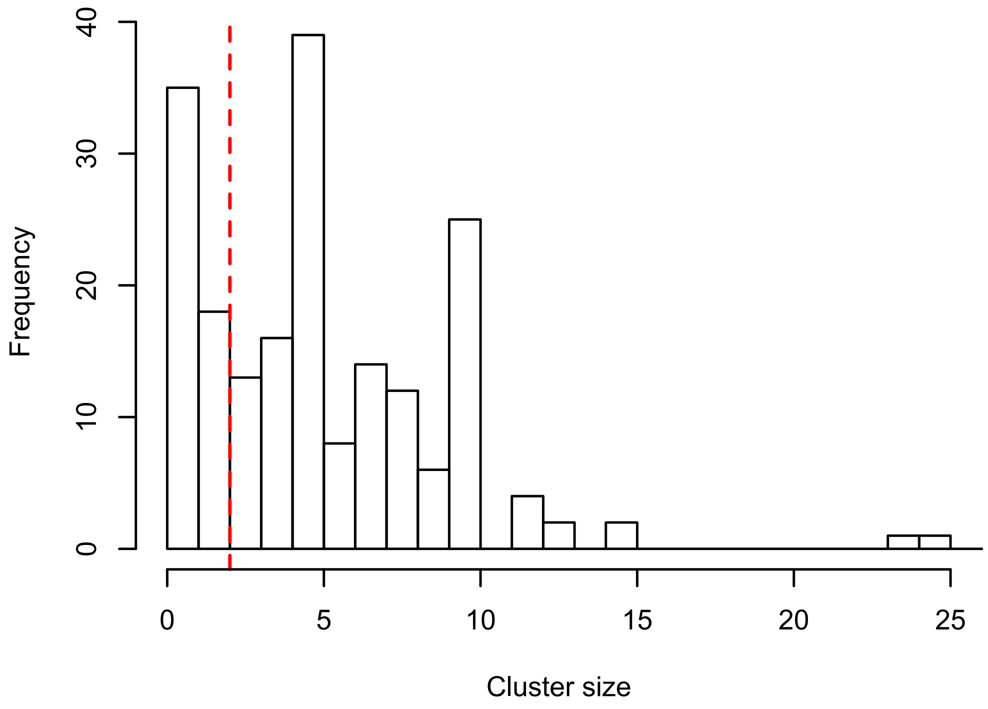


**Figure S2. Histogram of aggregated farm clusters based on a 4000m distance threshold.** Vertical line denotes a minimum cluster size of ≥3.

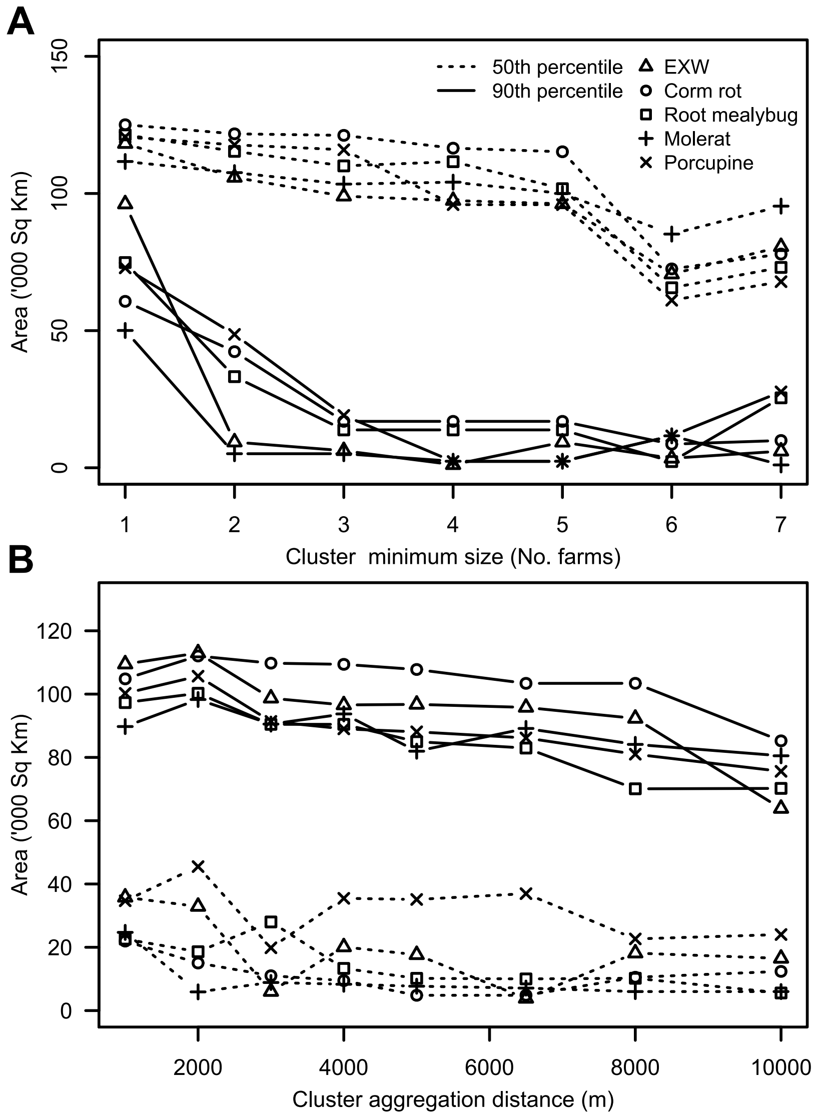


**Figure S3. Parameter sensitivity plots for niche envelope modelling.** Illustrating variability in predicted pest and pathogen distributions under A) a range of farm survey aggregation distances, and B) a range of farm survey minimum cluster sizes.

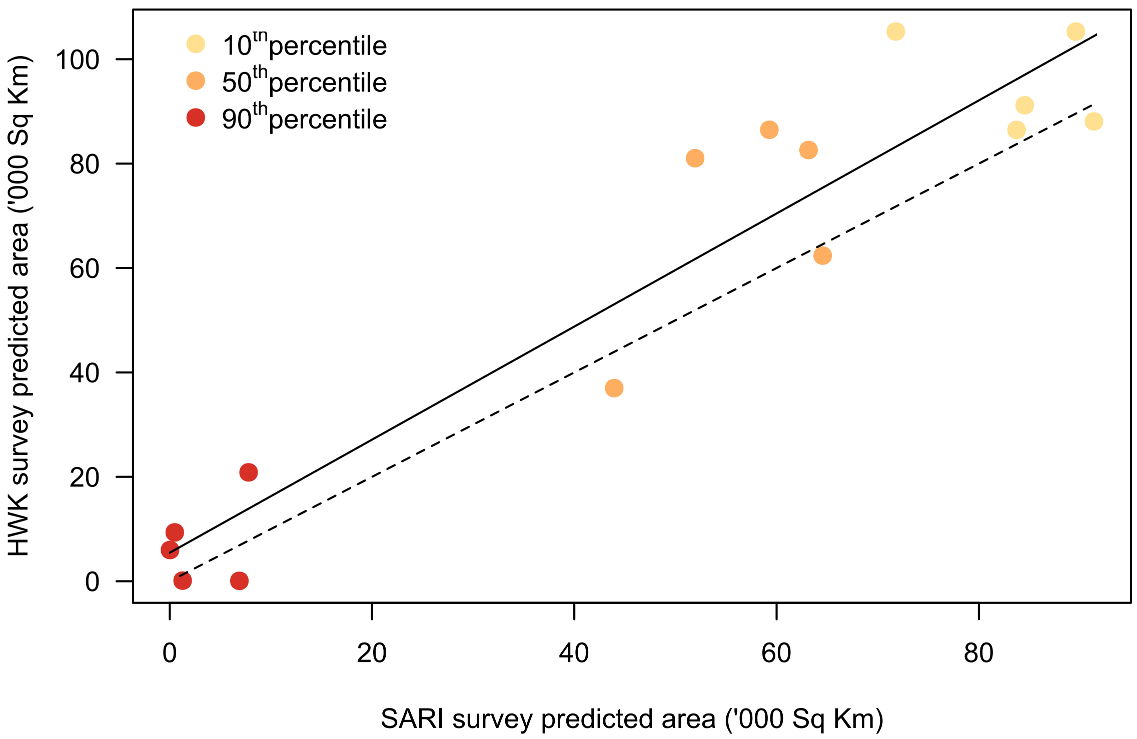


**Figure S4. Comparison of predicted area for each survey independently across three pest and pathogen incidence percentiles.** Dashed line indicates a 1:1 relationship.

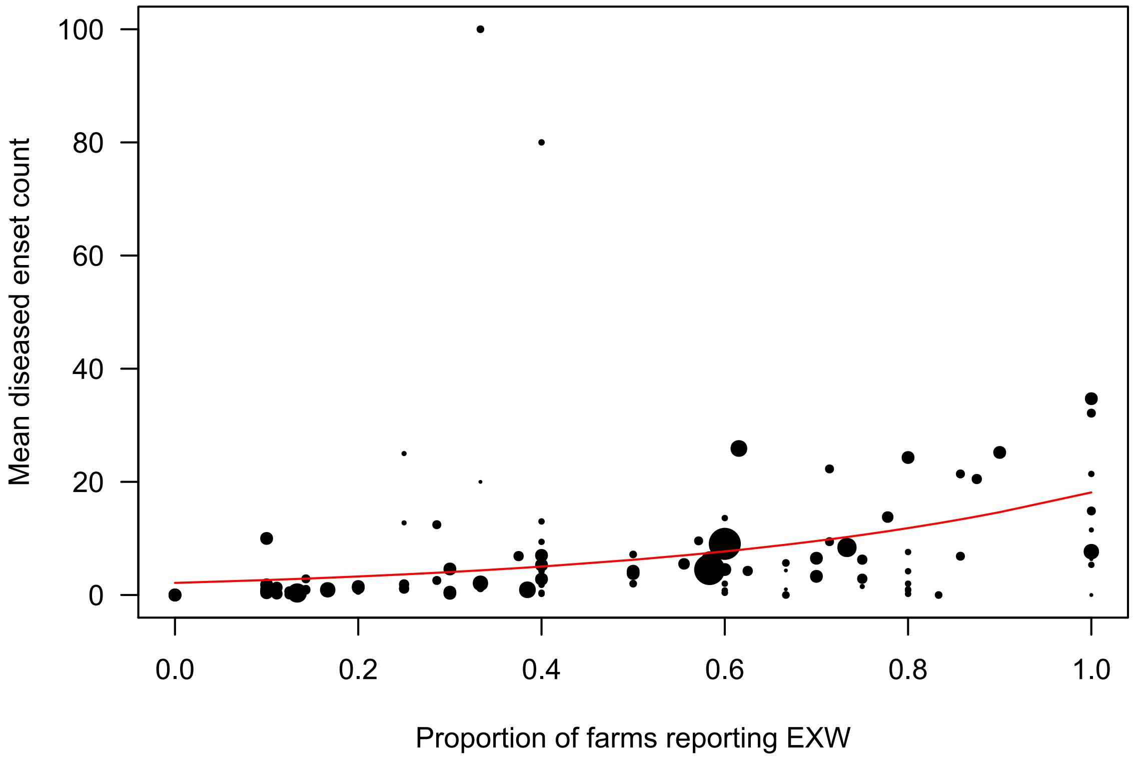


**Figure S5. Relationship between the proportion of farm clusters reporting EXW and the mean number of diseased enset recorded**, fitted with a quasipoisson generalized linear model and weighted by cluster sample size (range = 3-25).
